# Intraindividual longitudinal changes of the proteome reveal dynamic markers of biological aging

**DOI:** 10.64898/2026.10.01.755909

**Authors:** R. Lermer, X. Qian, S. Pusch, T. Schaefer, J. Adam, L. Li, M. von Scheidt, B. Linkohr, A. Peters, M. Mann, H. Schunkert, M. Heinig

## Abstract

Proteomic aging clocks have emerged as biomarkers of aging, but most rely on cross-sectional data. Here, we present a proteomic aging clock based on longitudinal measurements of liquid chromatography–mass spectrometry plasma proteomics at up to three time points in the population-based KORA cohort, comprising 9,105 samples from 4,100 individuals in the final dataset aged 25–88 years. The model predicts chronological age and yields a measure of proteomic age acceleration associated with lifestyle and clinical risk factors. Risk factor–stratified analyses reveal shared proteomic signatures among cardiometabolic exposures and distinct signatures for smoking and physical activity. Within individuals, dyslipidemia and impaired kidney function track transitions to increases in age acceleration, while higher age acceleration is associated with increased all-cause mortality. Mediation analysis suggests that risk factors are linked to mortality indirectly through age acceleration. Affected proteins and pathways highlight extracellular matrix remodeling, complement and coagulation pathways, and liver-derived plasma proteins as key contributors to aging. Mendelian randomization identifies proteins with evidence consistent with a causal role in aging-related traits. Together, these findings are consistent with biological aging being dynamically responsive to changes in modifiable risk factors.

## Introduction

Age is a major determinant of many chronic diseases^1^, yet chronological age incompletely captures interindividual heterogeneity in aging trajectories. The circulating proteome reflects systemic physiological processes and therefore provides a molecular window into biological aging^2^. Understanding how these molecular signatures evolve over time, and how they are shaped by modifiable risk factors, is central to identifying early changes that precede age-related disease.

Proteomic aging clocks (PACs) have emerged as integrative biomarkers of biological age, demonstrating associations with mortality and a broad range of clinical and lifestyle risk factors^2–6^, while organ-specific clocks highlight differential aging across tissues^3^. Most PACs have been developed using affinity-based plasma proteomics platforms^2,4,5^, which quantify predefined panels of proteins at high throughput but are inherently limited in proteome coverage. In contrast, liquid chromatography–mass spectrometry (LC-MS)–based proteomics enables unbiased, peptide-based quantification of thousands of plasma proteins with high analytical specificity^7^. As such, LC–MS provides complementary information to affinity-based approaches, with the potential to uncover novel molecular signatures of aging^8,9^, as first studies in small cohorts indicate^10^.

A key limitation of most existing PACs is their reliance on cross-sectional data, which do not capture within-individual changes in biological aging over time. Initial longitudinal studies have demonstrated the feasibility and prognostic value of repeated proteomic measurements. For example, analyses in the ARIC cohort linked proteomic age acceleration to mortality and cognitive outcomes across time points^11–13^. However, these studies either focused only on mortality^13^ or largely treated risk factors as baseline predictors and did not explicitly assess how changes in risk factor exposure relate to changes in biological aging.

Understanding whether within-individual changes of modifiable risk factors are reflected in corresponding changes in biological age is critical for advancing aging clocks from static risk predictors to dynamic biomarkers. Based on over 9,000 samples from a population-based cohort in Germany, analyzed with LC–MS at three time points, we developed a longitudinal proteomic aging clock. The model yields risk factor-specific signatures of key proteins and pathways contributing to aging, including dyslipidemia and kidney function, and predicts mortality. Integrating Mendelian randomization analyses, we further identify a subset of proteins with evidence consistent with a causal role in aging-related traits. Importantly, leveraging repeated measurements, we show that within-individual changes in proteomic age acceleration track changes in modifiable risk factors, including kidney function, BMI and dyslipidemia. Mediation analysis suggests that risk factors are linked to mortality indirectly through age acceleration. Together, these findings are consistent with biological aging being dynamically responsive to changes in modifiable risk factors.

## Results

### Training and performance

The KORA (Cooperative Health Research in the Region of Augsburg) cohort comprises longitudinal LC–MS plasma proteomics measured in 4,100 individuals at up to three time points (S4, F4 and FF4) approximately 7 years apart, spanning an age range of 25–88 years (Fig. 1a, Extended Data Fig. 1, Table 1). For this analysis a total of 9,105 samples were used.

**Fig. 1.**
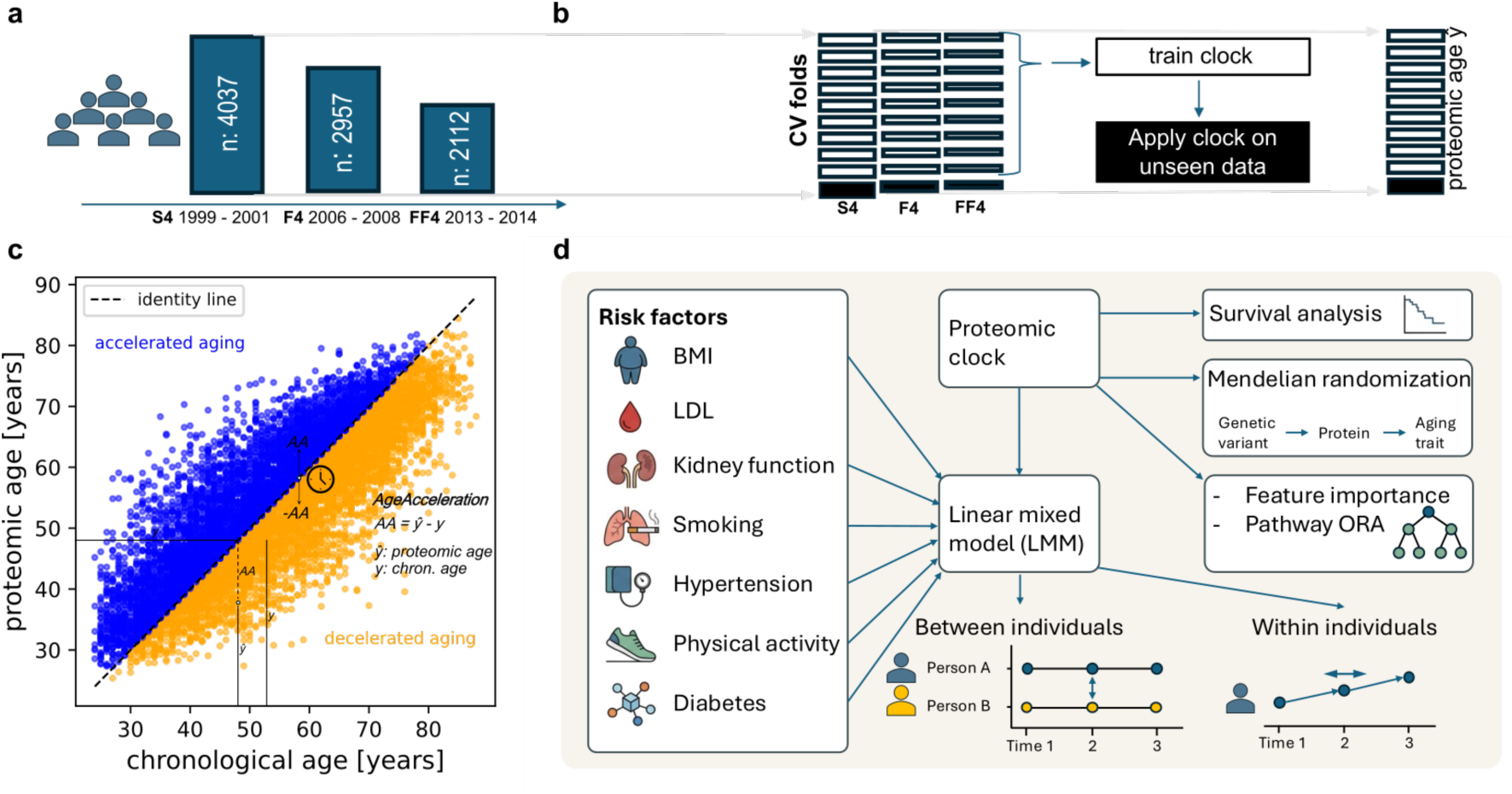
Overview. **a** KORA proteomics cohort: timepoints S4 and the two follow-up time points F4 and FF4 and their numbers of samples. **b** Cross validation over subjects based on the IDs in order to separate persons into different folds. **c** Scatterplot for estimated age illustrating the concept of age acceleration by formulas: Age acceleration is the difference between estimated (biological) and chronological age. **d** Visualization of the data analysis workflow. Age acceleration is linked to risk factors of lifestyle and environment and to blood biomarkers using linear mixed models assessing association of risk factors and age acceleration between individuals and associations of change of risk factor exposure and change in age acceleration over time within individuals. Subsequently the proteomic age is validated in a survival analysis. Also the indirect effects of risk factors via AA on mortality are assessed. Feature importance of proteins by SHAP values followed by overrepresentation analysis reveals pathways crucial for the process of aging. Mendelian randomization based on pQTLs and external GWAS data of aging associated traits pinpoints putative causal proteins.

**Table 1:** Table describing the cohort characteristics at each of the 3 timepoints. Detailed description of the variable definitions can be found in the methods.

| **Variable, unit** | **All** | **S4** | **F4** | **FF4** |
| --- | --- | --- | --- | --- |
| **Year** | 1999-2014 | 1999-2001 | 2006-2008 | 2013-2014 |
| **Final number of samples** | 9105 | 4037 | 2957 | 2112 |
| **Age mean, years** | 54.1 ± 14.1 | 49.2 ± 13.9 | 56.2 ± 13.3 | 60.3 ± 12.34 |
| **Age range, years** | 24 - 88 | 24 - 75 | 31 - 82 | 38 - 88 |
| **Sex (male)** | 48.9% | 49.3% | 48.10% | 49.10% |
| **BMI, mean** | 27.51 ± 4.82 | 27.24 ± 4.70 | 27.66 ± 4.82 | 27.81 ± 4.99 |
| **Diabetes** | 4.3% | 3.3% | 5.2% | 5.0% |
| **eGFR, mean** | 91.0 ± 18.5 | 95.4 ± 17.4 | 90.5 ± 18.3 | 83.2 ± 18.4 |
| **Ever Smoking** | 55.8% | 55.1% | 55.2% | 55.8% |
| **Hypertension** | 38.0% | 37.0% | 38.4% | 39.3% |
| **LDL > 130 mg/dl** | 54.1% | 54.1% | 53.7% | 52.4% |
| **Physical activity** | 52.2% | 48.3% | 54.1% | 56.9% |

To train the proteomic aging clock, we employed a multilayer perceptron with 10-fold cross-validation. To prevent information leakage across time points, folds were defined at the participant level, such that all samples from a given individual were assigned to the same fold (Fig. 1b). Hyperparameters were optimized by grid search (Extended Data Tables 1 and 2).

Model performance was evaluated by predicting chronological age. The final model achieved a mean absolute error (MAE) of 6.39 years and a Pearson correlation of 0.82 in cross-validation (Extended Data Table 3). Additionally, we assessed generalization between timepoints by training a clock on the S4 and FF4 subcohort and applying it on the F4 subcohort (Extended Data Fig. 2) and compared the performance to an elastic net model (Extended Data Fig. 3). Age acceleration (AA) was defined as the difference between predicted proteomic age and chronological age^14^ (Fig. 1c).

### Between-individual exposure to risk factors and proteomic age acceleration

Although the primary aim of this study was to investigate longitudinal changes in biological aging, we first sought to establish that the LC–MS–based proteomic clock captures biologically meaningful variation in aging-related risk factors between individuals. We therefore examined the association between AA with the modifiable risk factors BMI, diabetes, eGFR^3^, ever smoking, hypertension, physical activity^15^ (and subsequently with mortality^16^): For the risk factors we used linear mixed-effects models that decompose each risk factor into between- and within-individual components. This framework enables estimation of cross-sectional differences between individuals while accounting for repeated measurements over time (Fig. 2A). Risk factors were encoded as binary variables (e.g., LDL > 130), except for eGFR and BMI, which were modeled as a standardized continuous variable, scaled to its standard deviation of 18.52 units (ml/min/m^2^) and 4.82 (Kg/m^2^).

**Fig. 2.**
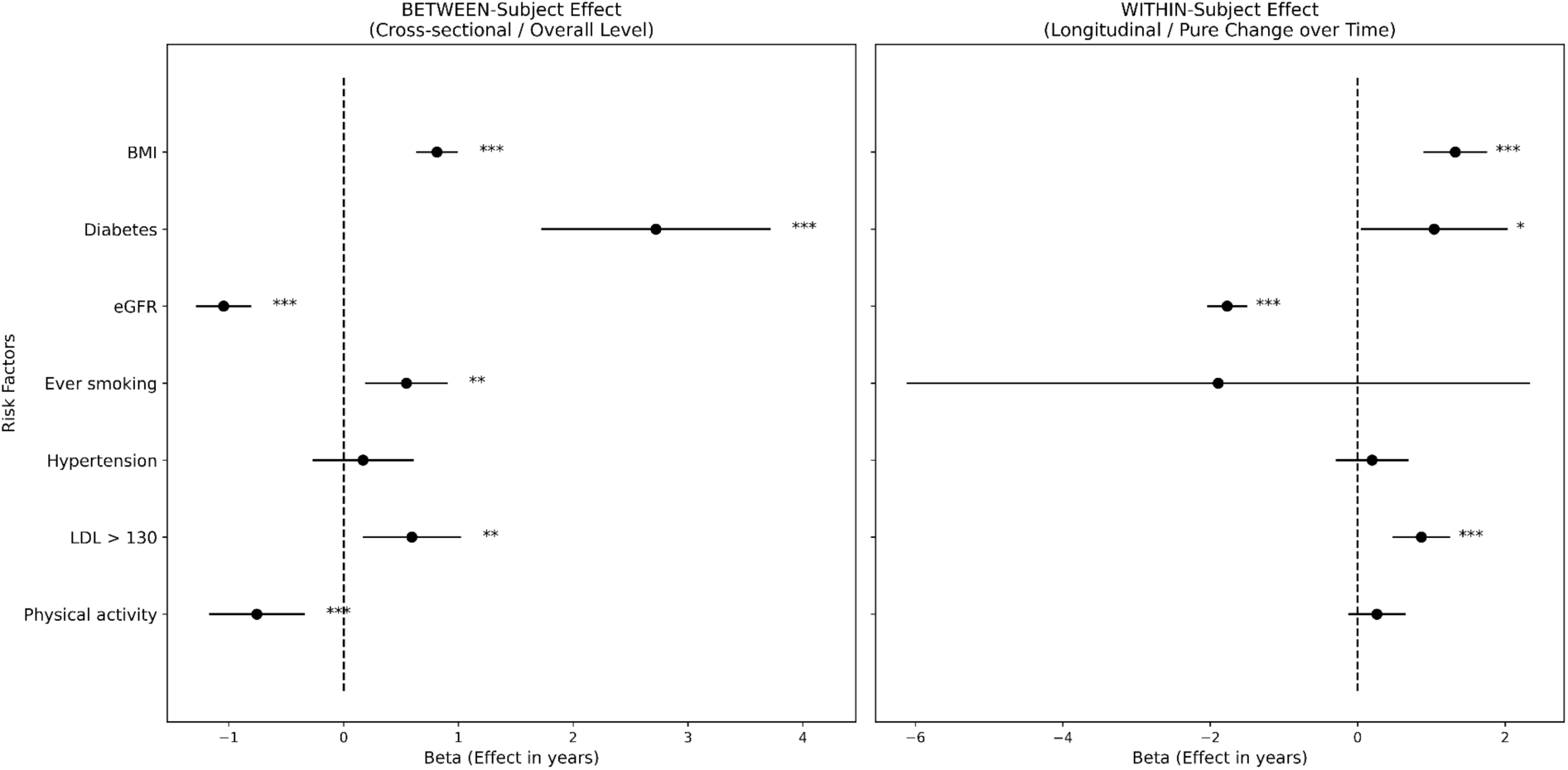
Forest plot. **a** Between-person effect sizes beta of linear mixed models that were fitted based on risk factors of lifestyle, environment and non-proteomic blood biomarkers, adjusted for age and age^2^. Beta for binary variables: difference of age acceleration between no and yes, for continuous variable(s) (BMI, eGFR): change of age acceleration for one standard deviation increase of the variable. **b** Within-person effect sizes beta show how changes in risk factor exposure within the same individual are associated with changes in age acceleration. Since the status of ever-smoking did only change for less than 10 persons in KORA, this risk factor did not entail a meaningful within effect.

Between-individual differences in cardiometabolic risk factors were strongly associated with proteomic age acceleration. Individuals with diabetes (β_between_ = 2.7 years, t = 5.3, P = 9.2 × 10^−8^) exhibited the largest increase in AA, corresponding to an effect size of approximately 2.7 years. Similarly, higher average exposure to BMI (β_between_ = 0.81 years, t = 8.7, P = 2.7 × 10^−18^), elevated LDL levels (>130 mg/dl) (β_between_ = 0.6 years, t = 2.7 × 10^−3^, P = 6.6 × 10^−3^), and smoking (β_between_ = 0.5 years, t = 3.0, P = 2.9 × 10^−3^) were all significantly associated with higher proteomic age acceleration.

In contrast, protective factors were associated with lower proteomic age acceleration. Higher average kidney function showed the strongest negative association (eGFR: β_between_ = −1.0 years per s.d., t = −8.5, P = 1.5 × 10^−17^), and higher levels of physical activity were also associated with reduced age acceleration (β_between_ = −0.8 years, t = 3.6, P = 3.6 × 10^−4^). These associations are consistent with previous reports from proteomic^5^ and epigenetic aging clocks^17,18^, supporting the robustness of the LC–MS–based proteomic clock.

### Longitudinal effects of risk factors

We next assessed whether temporal changes in risk factor exposure within individuals were associated with changes in proteomic age acceleration, using within-intraindividual components of the same linear mixed-effects models (Fig. 2B). The strongest effect was observed for eGFR, where one standard deviation higher than the mean within individuals was associated with an decrease of approximately 1.8 years in AA (β_within_ = −1.8 years per s.d., t = −12.6, P = 1.5 × 10^−36^). Notably, eGFR was derived from cystatin C and creatinine levels, which were not part of the proteomic input data, indicating that this relationship is not driven by direct model input features.

In contrast, increases in BMI (β_within_ = 1.3 years, t = 5.99, P = 2.09 × 10^−9^) in LDL levels (>130 mg/dl) were associated with higher age acceleration (β_within_ = 0.9 years, t = 4.3, P = 1.4 × 10^−5^), as were Diabetes (β_within_ = 1.0 years, t = 2.0, P = 4.2 × 10^−2^). No significant associations were detected for temporal changes of other risk factors, largely due to the fact that respective exposure was rather stable.

In addition, we tested whether risk factor exposure at a previous time point (t − 1) was predictive of AA at the following time point (t). In line with the analyses between and within individuals, the strongest positive effects were observed for BMI and diabetes, followed by smoking and elevated LDL (Extended Data Fig. 4). In addition, hypertension also showed a positive effect. Higher eGFR showed a strong negative association with future AA, whereas physical activity was not associated with future AA.

### Mortality prediction

Given that proteomic age acceleration tracked both prevalent risk factor burden and longitudinal changes in risk factor exposure within individuals, we next asked whether these aging-associated proteomic changes were also linked to long-term survival^5^. Within the KORA cohort, follow-up of up to 22 years allowed to demonstrate strong associations between AA and all-cause mortality. Specifically, individuals in the highest and lowest deciles of AA (stratified by chronological age) showed a clear separation in survival probabilities (Fig. 3a; log-rank test, P = 8.66 × 10^−11^), also when analysing time points separately (log-rank test, S4: P = 2.3 × 10^−6^, F4: P = 2.4 × 10^−3^, FF4: P = 1.5 × 10^−3^).

**Fig. 3.**
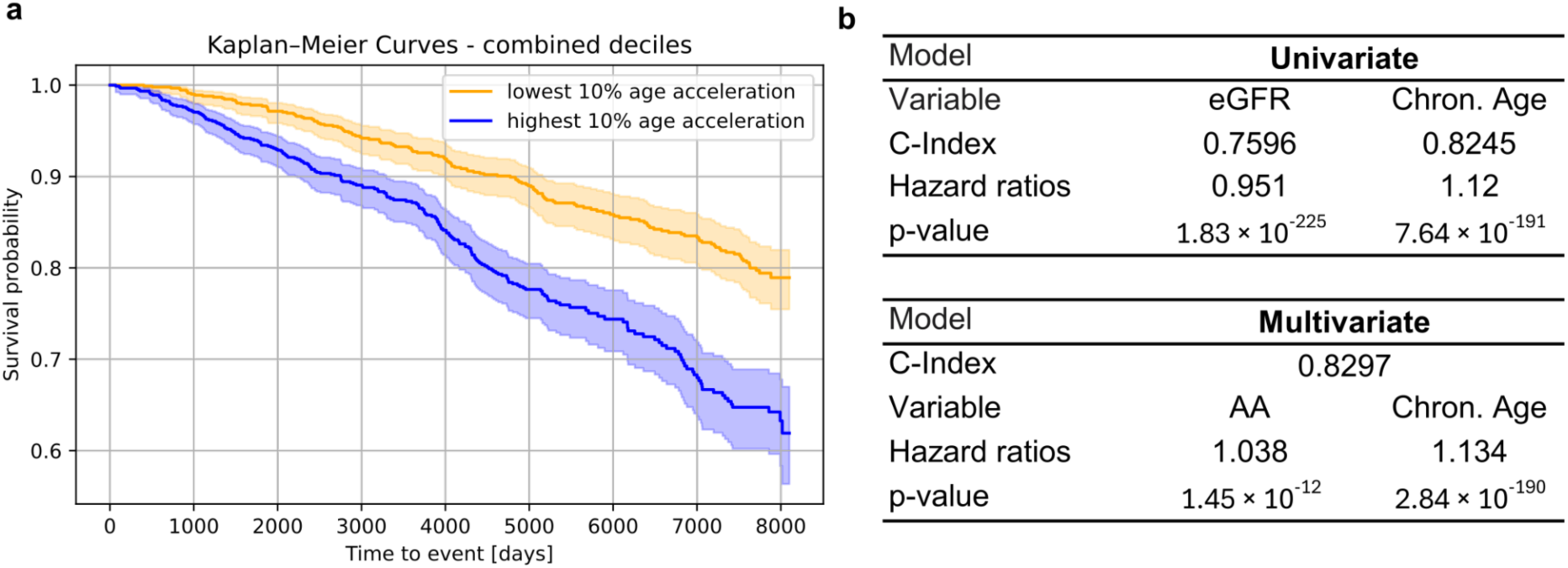
Survival analysis. **a** Kaplan-Meier plot of two groups: top and lowest 10 percent of age acceleration stratified by deciles of chronological age. **b** Results of survival analysis for univariate models with only chronological age and eGFR as predictors and a multivariate model with the predictors chronological age and age acceleration.

In Cox proportional hazards models, inclusion of AA alongside chronological age improved mortality prediction compared to age alone (C-index 0.830 vs. 0.825), with AA contributing independently (Wald test, P = 1.45 × 10^−12^; Fig. 3b). This association remained significant after adjustment for kidney function (eGFR) (Extended Data Table 4, Extended Data Fig. 5). Together, these findings suggest that the proteomic changes associated with accelerated biological aging reflect physiologically relevant processes linked to long-term health outcomes.

### Mediation analysis of risk factors, age acceleration and mortality

Given that risk factor exposure is associated with AA and AA in turn is predictive for mortality, we tested whether AA might mediate part of the association of risk factors with mortality. We selected risk factors associated with mortality in the S4 subcohort (Extended data Table 5) for this analysis. We found significant indirect effects for all tested risk factors except LDL > 130, using G-Computation with bootstrapping and Monte-Carlo-Simulations in the S4 subcohort (Fig. 4). According to the mediation model, the largest indirect effect was observed for diabetes, for which the pathway through AA corresponded to an increase in mortality risk of 2.01% (1.12% - 3.31%) of a total effect of (10.33%) (proportion mediated: 0.195). Indirect effects leading to increased risk were also observed for BMI (indirect: 0.47% (0.28% - 0.67%), total: 1.72%, proportion mediated: 0.27), ever smoking (indirect: 0.21 (0.011 - 0.45), total: 5.21, proportion mediated: 0.041) and hypertension (indirect: 0.35 (0.12 - 0.65), total: 2.10, proportion mediated: 0.168). Reduction in risk was attributed to indirect effects of eGFR (indirect: −0.41% (−0.73% - −0.23%), total: −1.66%, proportion mediated: 0.25) and physical activity (indirect: −0.16 (−0.36 - −0.0036), total: −4.98, proportion mediated: 0.033).

**Fig. 4.**
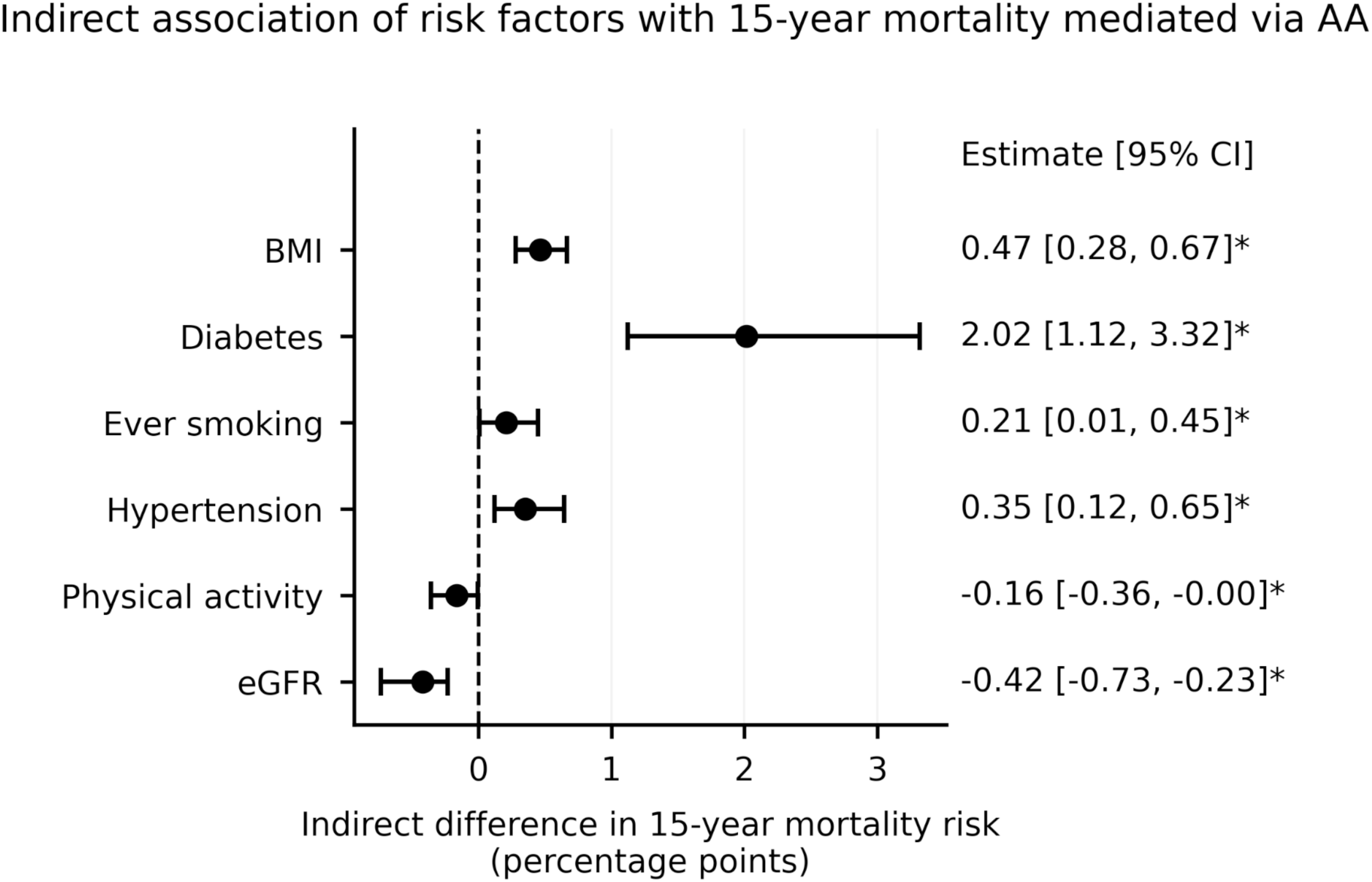
Forest plot indirect effects of risk factor exposure in S4 on mortality mediated over age acceleration. Indirect effects of risk factors (percentage points of elevated mortality risk) through age acceleration show that most of the risk factors are associated with a higher age acceleration which can be associated with a higher mortality risk. These results are consistent with a causal pathway from risk factor exposure through AA to mortality risk.

### Proteins associated with fast and slow aging

To gain insight into the molecular drivers of proteomic age prediction, we quantified protein importance using SHAP values (Fig. 1f). Ranking proteins by mean absolute SHAP values identified those contributing most strongly to biological age estimation.

The top 25 proteins (Fig. 5a) included LUM, ALS and AFAM as the most influential features. Most of these proteins have previously been linked to aging, age-related traits or mortality (Extended Data Table 6), indicating that the model captures established molecular signatures of aging. Notably, overlap with affinity-based proteomic clocks was limited: none of the top features were shared with the ProtAge clock^5^, and only APOB overlapped with the SomaScan-based model by Lehallier et al.^2^, consistent with differences in proteome coverage across platforms^8^.

**Fig. 5.**
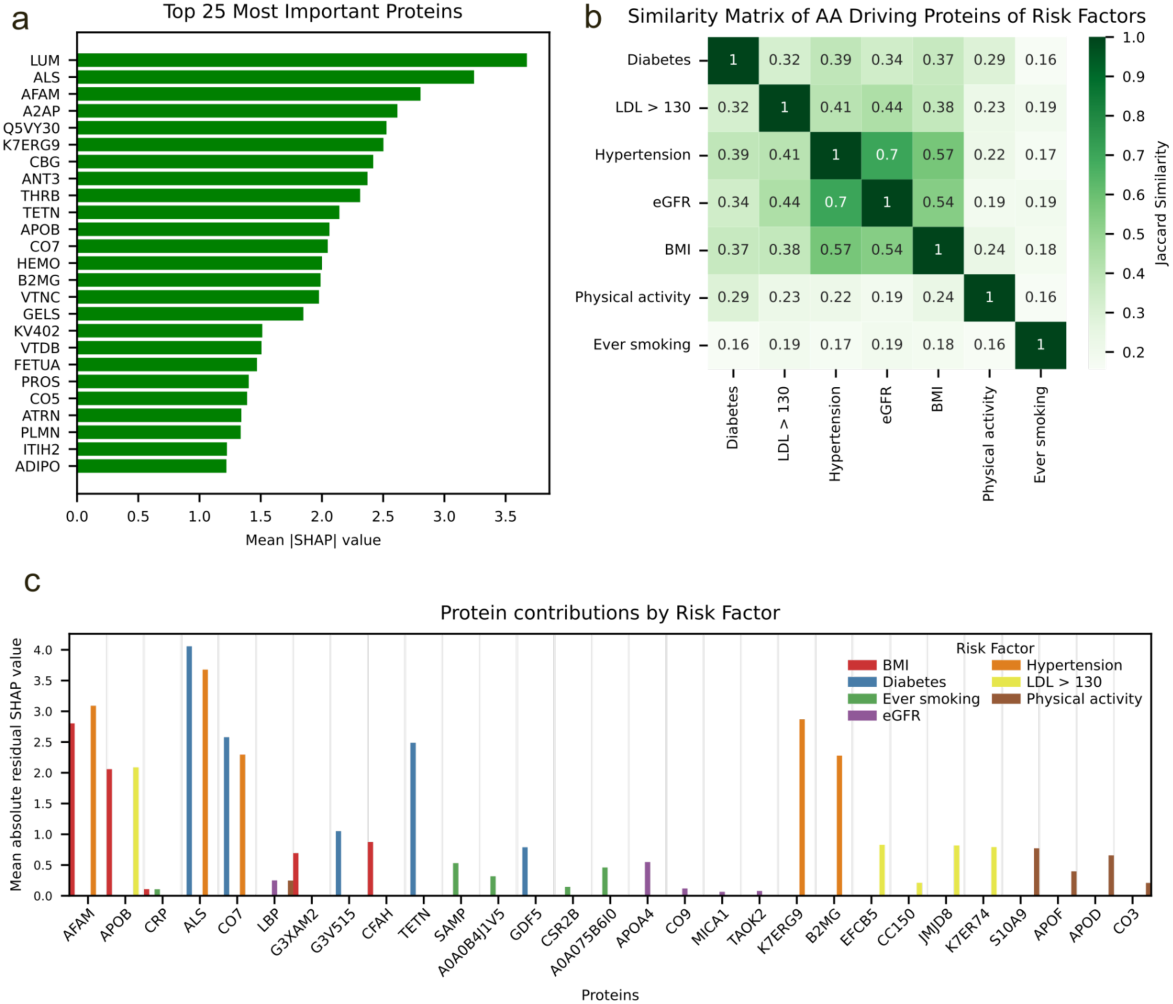
Feature importance. **a** Top 25 proteins’ mean absolute SHAP values. **b** Similarity of proteomic profiles by risk factor: Each risk factor gained a different proteomic profile driving the age acceleration (see methods). We calculated the similarity measures (Jaccard indices) between those profiles. **c** Top five proteins for each risk factor driving age acceleration (mean residual SHAP values)

Pathway enrichment analysis of the top-ranked proteins revealed significant overrepresentation of complement, coagulation and xenobiotic metabolism pathways (including VTN, RBP4, PROS1, FETUB, PLG, FBLN1, APOE, HRG, SERPINA6 and PTGDS), which predominantly represent liver-derived plasma proteins, pointing to alterations in the circulating liver secretome (Extended Data Table 7). These findings highlight coordinated changes in hepatic, inflammatory and thrombotic processes that have previously been implicated in aging^2,19,20^.

We next examined whether the molecular drivers of age acceleration differ across risk factor exposures. Using residual SHAP values (see methods) to isolate protein contributions to age acceleration, we identified risk factor-specific proteomic signatures (Fig. 5c). While several proteins were shared across cardiometabolic risk factors, particularly diabetes, hypertension and BMI, most top-ranked proteins were specific to individual exposures.

To quantify these relationships, we compared proteomic signatures using the Jaccard index (Fig. 5b). Cardiometabolic risk factors (BMI, hypertension and reduced eGFR) showed substantial overlap (Jaccard index > 0.5). Dyslipidemia and Diabetes also showed moderate overlap with those and to each other (> 0.3). In contrast, smoking and physical activity exhibited distinct proteomic signatures with minimal overlap with other risk factors.

Overrepresentation analysis (ORA) of residual SHAP values for each risk factor identified distinct pathway enrichments (Extended Data Table 8). For hypertension, xenobiotic metabolism and angiogenesis were enriched, while for eGFR, xenobiotic metabolism was identified. For dyslipidemia (LDL > 130), the complement pathway was significantly enriched. Collectively, these results indicate that accelerated aging related to specific risk factor exposures is associated with coordinated alterations in hepatically derived plasma proteins, vascular extracellular matrix processes, and thrombo-inflammatory pathways. Specifically, cardiometabolic risk factors share common proteomic signatures linked to immune and thrombo-inflammatory pathways, whereas smoking and physical activity are characterized by distinct molecular profiles. These findings link accelerated biological aging with exposure-specific molecular processes.

### Inference of proteins causally relevant for aging acceleration

Mendelian randomization (MR) analysis (Fig. 6) was carried out to link aging-related proteins (pQTLs, Supplementary table 9) genetically with aging related traits (frailty, longevity, parental lifespan, healthspan, intrinsic epigenetic age acceleration, multivariate aging). Among the 842 proteins analyzed, 28 showed evidence consistent with a causal relationship between protein expression and at least one aging-related trait (Supplementary Table 10). Notably, 14 of these proteins were among the top 20% most important features for proteomic age prediction (Fig. 6), representing a significant enrichment compared to the expected overlap of 5.6 proteins under a random model (OR = 4.3, P = 2.8 × 10⁻^4^, Fisher’s exact test). These findings suggest that proteins driving the proteomic aging signal are not only associated with aging phenotypes but are enriched for putative causal mediators of aging-related processes, such as APOE (Fig. 6) or CO4A.

**Fig. 6.**
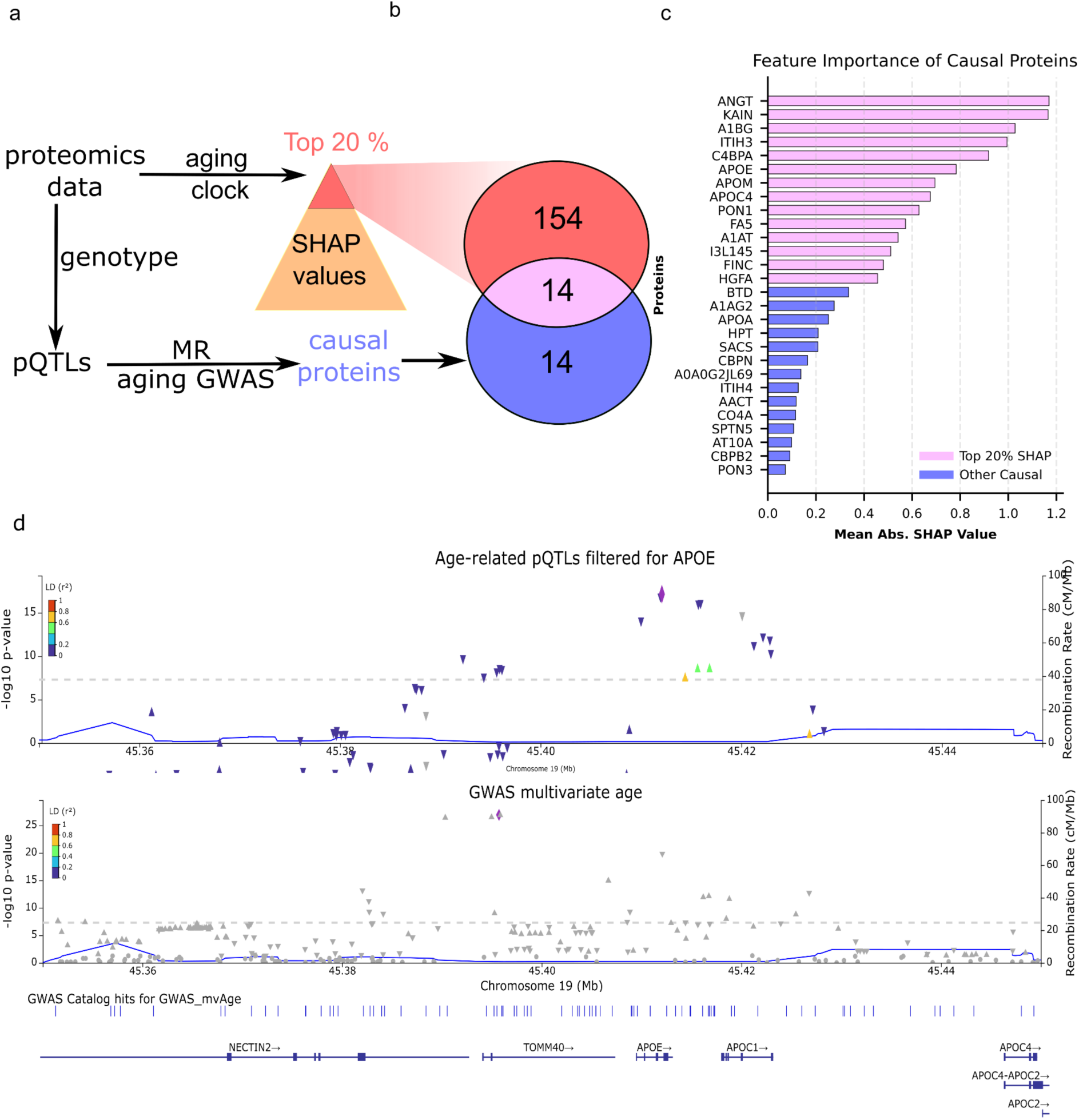
Mendelian randomization of proteins and aging related traits. **a** Schema for the workflow to identify proteins with high feature importance and causal for aging **b** Venn diagram of those proteins **c** Barplot showing feature importance of causal proteins **d** Zoomed manhattan plots in the region around APOE for pQTL data filtered for APOE and GWAS data about multivariate aging

## Discussion

In this study, we developed a mass spectrometry–based proteomic aging clock in a population-based cohort with longitudinal sampling across up to three time points. In line with previous studies^5,17,18^, we observed that prevalent risk factors accelerate proteomic age, which also associates with mortality. Indeed, the plasma proteome may mediate some of the detrimental effects of modifiable risk factors, as supported by Mendelian randomization analyses identifying a subset of proteins as putative causal mediators of aging-related traits. A central advance of this work is the explicit modeling of both between-individual differences and intraindividual changes in risk factor exposure using a unified longitudinal framework. While previous longitudinal studies revealed that risk factors, including impaired kidney function, at baseline can predict subsequent age acceleration^11^ they have largely treated risk factors as static predictors. By contrast, our approach disentangles cross-sectional from longitudinal effects and demonstrates that temporal changes in risk factor exposure within individuals are accompanied by corresponding changes in proteomic age acceleration. Because the within-individual estimates compare each participant with themselves over time, they are not confounded by characteristics that remain stable within a person, such as genetic background, early-life exposures or socioeconomic position, which limit the interpretation of cross-sectional associations.

Specifically, we show that transitions toward diabetes, dyslipidemia, and BMI within individuals were associated with increases in proteomic age acceleration, whereas impaired kidney function is associated with elevated age acceleration. These findings extend prior work by demonstrating that biological aging not only reflects past and current risk factor exposure, but also ongoing changes in physiological state over time. Importantly, the consistency between between-individual and within-individual associations, as well as the association between past risk factor exposure and current biological age suggests that risk factors linked to accelerated aging at the population level also contribute to dynamic changes in biological aging trajectories within individuals.

The association between changes in kidney function and proteomic age highlights the close link between renal and systemic aging. Previous studies have linked eGFR to aging clocks and age-related disease risk^3,5^ and longitudinal analyses have associated baseline kidney function with future age acceleration^11^. Our results extend these findings by demonstrating that intra-individual changes in kidney function are accompanied by corresponding changes in biological aging, supporting the concept that renal and systemic aging are tightly coupled processes.

Risk factor-specific analyses revealed both shared and distinct molecular signatures of aging. Cardiometabolic risk factors, including diabetes, hypertension, dyslipidemia and reduced kidney function, showed overlapping proteomic signatures enriched for complement, coagulation and liver-associated pathways, whereas smoking and physical activity were characterized by more distinct profiles. These findings suggest that accelerated aging reflects a combination of shared cardiometabolic processes and exposure-specific molecular responses.

The observed pathways reflected by plasma proteomics point to an integrated liver–vascular–inflammatory axis underlying systemic aging. Alterations in liver-derived plasma proteins likely reflect changes in systemic metabolic and secretory function, while extracellular matrix remodeling and vascular dysfunction are hallmarks of vascular aging^21^. In parallel, chronic low-grade inflammation promotes complement activation and pro-thrombotic states that contribute to age-related disease^22,23^. Together, these processes support a model in which thrombo-inflammatory and metabolic pathways jointly drive systemic biological aging.

Our findings have important implications for aging research and prevention. The association between changes in risk factor exposure and corresponding changes in proteomic age suggests that biological aging is not fixed but can be modified, at least in part, through changes in modifiable risk factors. Given the link between age acceleration and mortality risk, these results imply that improvements in risk factor profiles may translate into gains in healthspan and potentially lifespan. Indeed, the results of the mediation analysis support this idea, indicating that indirect effects of risk factors on mortality might be mediated via AA with the largest indirect effect observed for Diabetes and BMI and eGFR the largest proportions mediated. More broadly, the identification of risk factor-specific proteomic signatures, together with evidence for causal involvement of key proteins, highlights potential molecular targets for intervention and provides a framework for interpreting aging clocks across different biological modalities.

At the molecular level, proteins contributing to age prediction were enriched for extracellular matrix components, complement and coagulation pathways, and liver-derived plasma proteins, consistent with established signatures of systemic aging. Despite limited overlap with affinity-based proteomic clocks likely reflecting differences in protein coverage across technologies^8^, our results converge on similar biological processes, providing independent validation of these pathways. Importantly, Mendelian randomization analyses linking genetically predicted protein levels to aging-related traits further supported the biological relevance of these proteins. Proteins identified as important for age prediction were significantly enriched for putative causal mediators of aging-related traits, suggesting that the proteomic aging signal captures molecular pathways that are not only associated with, but may contribute to, aging-related processes.

Several limitations should be considered. First, the predictive accuracy of the model was lower than that reported for some larger proteomic or DNA methylation–based clocks^2,4,5^, potentially reflecting the sample size and the focus on longitudinal generalizability. Second, the cohort is limited to a Western-European population. An independent longitudinal proteomic cohort with repeated LC-MS measurements was not available for replication of the within-individual analyses and will be important for future validation. Third, plasma proteomics captures only a subset of biological processes relevant to aging. Fourth, although the within-individual analyses remove confounding by stable characteristics, they do not account for time-varying confounders, and the mediation analysis relies on assumptions that cannot be verified in observational data.

In conclusion, this study provides a longitudinal perspective on molecular aging by demonstrating that both differences between individuals and changes within individuals in modifiable risk factors are reflected in proteomic age acceleration. The integration of longitudinal modeling with genetic evidence further suggests that key proteins contributing to the proteomic aging signal may play a causal role in aging-related processes. These findings support the concept that biological aging is dynamic and potentially modifiable, and establish mass spectrometry–based proteomics as a powerful platform for studying aging trajectories and their underlying molecular mechanisms.

## Methods

### Cohort

We used plasma proteomic mass-spec data from the population-based KORA (Cooperative Health Research in the Region of Augsburg) S4/F4/FF4 study.^24^ The study consists of three timepoints called S4, F4 and FF4 with approximately seven years between the timepoints. They were sampled in the years 1999 - 2001 (S4), 2006 - 2008 (F4) and 2013 - 2014 (FF4). The age range of the subjects is 24 up to 88 years over all the three timepoints and initially 24 to 75 years at timepoint S4. Details are shown in Table 1. Standardized examinations were conducted by trained staff and included a face-to-face interview on health status and life-style factors, blood pressure measurements, anthropometry, and blood draw. Details on the study characteristics are shown in Table 1. Mortality was assessed between baseline and the end of the follow-up on Dec. 31, 2021 using death certificates from local health authorities and the vital status from population registries. The follow-up time was calculated from baseline examination until the date of death for deceased and for living persons until the date of last contact.

We define the variables used in this analysis as follows: Age refers to the age at the day of examination. Diabetes status is encoded as 1 if participants had a HbA1c value larger than 48 and 0 otherwise. If the diabetes status was unclear, they were excluded from the analysis. The estimated Glomerular Filtration Rate (eGFR) (ml/min/1.73m^2^) was calculated based on serum creatinine and cysteine concentrations using the Chronic Kidney Disease Epidemiology Collaboration formula. Physical activity was assessed by self declaration in a face-to-face interview and was categorized as active or inactive based on the frequency and duration of weekly exercise in summer and winter. Smoking status was classified as current smoker, ex-smoker, or never-smoker. BMI was calculated as the body mass divided by the squared body height in Kg/m^2^. Low-density lipoprotein (LDL) cholesterol levels were measured by enzymatic photometric assays in S4 and by enzymatic colorimetric Flex assays in F4 and FF4.

### LC–MS/MS proteomics analysis

A non-targeted liquid chromatography–tandem mass spectrometry (LC–MS/MS) workflow was used to quantify relative protein abundances. Blood plasma samples were collected at each examination time point and stored at −80 °C until analysis. Samples from all three time points (n = 9,201, initially) were analyzed in a single batch between December 2021 and May 2022 to minimize technical variability.

Sample preparation largely followed the protocol described by Geyer et al^25^ with the modification that a detergent-free lysis buffer was used instead of an SDC-based buffer. Briefly, 1 µL of plasma was subjected to simultaneous denaturation and alkylation at 95 °C, followed by enzymatic digestion with trypsin and LysC at an enzyme-to-protein ratio of 1:100 at 37 °C for 4 hours, yielding peptides for LC–MS/MS analysis. Peptides were separated online by nanoflow reversed-phase liquid chromatography using an Evosep One system coupled to a timsTOF Pro2 mass spectrometer. We used the dataset from Niu et al^26^ with the same quality control version, incorporating one additional step: We removed samples with more than 30% missing before the normalization. This reduced the number of samples from 9,201 to 9,105. In the final dataset the number of proteins was 842. Feature wise quantile normalization as a final step before the data was used for the training.

### Training of the proteomic clock model

We aimed to estimate a proteomic age for every person in the cohort. Before the training, the normalized and quality controlled proteomics dataset was quantile-transformed to make it better aligned for a regression model. It has been shown before that protein abundance over lifetime shows non-linear patterns.^2^ Therefore, we employed neural networks for our proteomic clock, as they can exploit non-linear pattern.^27^ We set up a multi layer perceptron (MLP) to predict chronological age based on the proteomics data. The model was implemented in PyTorch.^28^ We performed a sweep tuning on a 80/20 split to find the best hyperparameter for the network. The hyperparameters can be found in extended data table 1 and the optimal hyperparameters in extended data table 2. Using these hyperparameters, we performed a ten fold cross validation which places all samples from one person into one fold. In addition, we compared the performance of the neural network to an elastic net model, using the same cross validation approach (Extended Data Fig. 3).

### Estimation of proteomic age and age acceleration

The optimized hyperparameters were used to train the final model within a ten-fold cross-validation framework to estimate proteomic age for each individual. To prevent data leakage across time points, cross-validation folds were defined at the participant level, such that all samples from the same individual were assigned to the same fold. This ensured that no samples from the same person were present in both training and test sets. Moreover, it ensured that biological age prediction for an individual is not based on a model that was trained on this person’s data. Proteomic age acceleration was defined as the difference between predicted proteomic age and chronological age.^29^

### Analysis of risk factors (between individuals)

To assess between-individual associations of proteomic age acceleration and lifestyle, environmental, and clinical risk factors, we employed linear mixed-effects models that account for repeated measurements within individuals. This framework enables the simultaneous estimation of between-individual differences and within-individual changes over time.

For each risk factor, values were decomposed into a between-individual component, defined as the individual-specific mean across all available time points, and a within-individual component, defined as the deviation from this mean at each time point. The between-individual component captures stable differences in exposure levels across individuals, independent of temporal variation.

For each risk factor, we fitted a linear mixed-effects model of the form

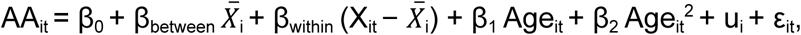

where AA_it_ denotes proteomic age acceleration for individual i at time point t, X_it_ the observed value of the respective risk factor, *X̄*_i_ the individual-specific mean of that risk factor across all available time points, u_i_ the subject-specific random intercept, and ε_it_ the residual error term. In this parameterization, β_between_ captures the between-individual association and β_within_ captures the within-individual association. The random intercept for each individual was included to account for within-subject correlation across repeated measurements. Chronological age and age² were included as fixed effects to account for non-linear age dependencies.

Regression coefficients for the between-individual components, along with their 95% confidence intervals, were used to quantify associations between average exposure levels and proteomic age acceleration and to generate the forest plot (Fig. 2).

### Analysis of risk factors (within-person effects)

Within-individual associations between changes in risk factor exposure and proteomic age acceleration were assessed using the within-individual component of each risk factor, defined as the deviation of the observed value at a given time point from the individual-specific mean across all available time points (X_it_ − *X̄*_i_). In the mixed-effects model above, the coefficient β_within_ therefore quantifies the extent to which temporal changes in risk factor exposure within an individual are associated with corresponding changes in proteomic age acceleration, independent of stable between-individual differences.

### Analysis of risk factors (between time point effects)

To assess between-timepoint associations of proteomic age acceleration and lifestyle, environmental, and clinical risk factors, we employed linear mixed-effects models that account for repeated measurements within individuals (“Lagged LMM”). For each risk factor X, we fitted a linear mixed-effects model of the form

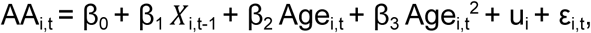

using age acceleration AA_i,t_ from F4 (t = 2) and FF4 (t = 3) paired with risk factor data X_i,t-1_ from S4 (t = 1) and F4 (t = 2). The variable u_i_ denotes the subject-specific random intercept, and ε_i,t_ denotes the residual error term.

### Survival analysis

To evaluate the association between proteomic age acceleration and all-cause mortality, Cox proportional hazards models^30^ were fitted using the lifelines python library.^31^ To account for the repeated measurements in the longitudinal data, we used the robust sandwich estimator of the variance of the effect size estimates.^32^ First, two univariate models were fitted using chronological age and eGFR as predictors. The model using chronological age as predictor serves as the null model, while the model based on eGFR serves as a second baseline for comparison. A multivariable model including chronological age and proteomic age acceleration was then fitted to assess whether age acceleration improved mortality prediction. Model performance was compared using the concordance index (C-index,^33^ and hazard ratios with corresponding p-values were reported.

For visualization, Kaplan–Meier survival curves were generated. To account for regression-to-the-mean effects across chronological age, participants were stratified into deciles of chronological age. Within each decile, individuals in the top and bottom 10% of age acceleration were identified and pooled across deciles to form two groups. Kaplan–Meier curves were compared using a log-rank test. To assess the robustness of the results, we also applied the log-rank test separately for each time point.

### Mediation analysis of risk factors, age acceleration and mortality

We aimed to test whether the association of a risk factor with age acceleration (AA) leads to indirect association with 15-year mortality risk through AA. We used G-Computation and Monte Carlo simulations to estimate the 15-year mortality risk in the S4 subcohort to do so. Participants (12) that were censored alive before the end of the 15-year horizon, were removed from the analysis. Continuous risk factors were z-standardized and the chronological age was mean centered.

Firstly, we tested for each risk factor if it is associated with mortality using logistic regression models for each risk factor and adjusted them for centered age and centered age squared. These models were fitted on the S4 subcohort using 15 years of follow up time to estimate the absolute probability of death in 15 years of follow up time.

For the G-Computation we built several parametric models: The “mediator model” was estimated following this formula:

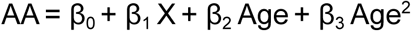

The “outcome model” was estimated following this formula:

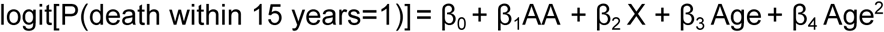

We used the python package pygformula to do the g-formula computation. Because pygformula requires time-indexed data, we represented each S4 observation by two technical simulation steps. At t0, the observed S4 values initialized the simulation, whereas at t1, age acceleration was modeled and the binary 15-year mortality outcome was evaluated.

Based on those we build three risk scenarios: R00 = Risk(X=0, M(0)), R10 = Risk(X=1, M(0)), R11 = Risk(X=1, M(1)). X=0 means that the risk factor is set to 0 or if it is a continuous risk factor set to the mean. Whereas X = 1 means the risk factor is set to 1 or for continuous variables one standard deviation above the mean. M(0) is the AA simulated by using the mediator model with X = 0 and M(1) accordingly for X = 1. The estimated risks represent the absolute probability of death within 15 years under each simulated intervention scenario. Risk differences were calculated on this absolute probability scale and multiplied by 100 to express them as percentage-point differences in 15-year mortality risk. For example, a risk difference of 0.02 corresponds to an absolute increase of 2 percentage points.

The indirect effect was calculated as R11−R10 and expressed as an absolute difference in 15-year mortality risk. The total effect was calculated as R11−R00. Pygformula estimates for every bootstrap the risks. We used 2000 bootstrap samples to calculate the percentile bootstrap confidence intervals for the simulated indirect effects ranging from the 2.5th to the 97.5th percentile. We considered the indirect effect as significant if zero was not included within the confidence interval.

A causal interpretation of these total and indirect effects requires the standard identification assumptions for mediation analysis, including consistency and positivity, no unmeasured confounding of the exposure–mediator, exposure–outcome, and mediator–outcome relationships conditional on the included covariates, and correct specification of the mediator and outcome models. In particular, the interpretation further assumes the absence of mediator–outcome confounders that are themselves affected by the exposure. As these assumptions cannot be fully verified from observational data, we interpret the estimated indirect effects as evidence consistent with a pathway through AA rather than as definitive evidence of causal mediation.

### Feature importance analysis

Feature importance for individual proteins was quantified using mean absolute SHAP (SHapley Additive exPlanations) values.^34^ To contextualize the 25 top ranked proteins in the literature, a PubMed search was performed for associations with aging, age-related diseases, or mortality. The results are summarized in Extended Data Table 1.

To identify proteins contributing specifically to age acceleration within risk factor subgroups, we leveraged the additive structure of SHAP values. For each sample, SHAP values sum to the difference between predicted age and the model base value (expected prediction). Because the base value corresponds to the mean chronological age, SHAP values primarily explain chronological age prediction. To focus on contributions to age acceleration, we computed residual SHAP values by removing the component attributable to chronological age. Assuming an equal expected contribution of all proteins to the chronological age y component, we distributed chronological age uniformly across all proteins and subtracted this expected contribution from each SHAP value. Specifically, for each sample i and protein j, residual SHAP values were computed as

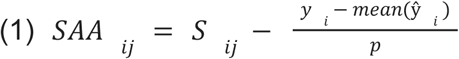

where S_ij_ denotes the SHAP value for protein j in sample i, y_i_ the chronological age, ŷ*_i_* the proteomic age and p the number of proteins. This transformation isolates protein contributions related to age acceleration. Risk factor–specific protein signatures were then identified as follows. For binary risk factors, individuals were stratified by exposure and protein-wise differences in residual SHAP values were tested using two-sample t-tests. For continuous risk factors (eGFR), residual SHAP values were regressed on the risk factor. Resulting p-values were adjusted using the Benjamini–Hochberg procedure. Proteins with adjusted p-values < 0.05 were considered part of the proteomic signature of the respective risk factor and ranked by mean absolute residual SHAP value. Similarity between risk factor–specific protein signatures was quantified using the Jaccard index and visualized as a similarity matrix (Fig. 5b).

### Gene set Over-Representation Analysis (ORA)

Gene set enrichment analyses were performed to identify biological pathways associated with proteins driving age prediction and risk factor-specific aging signatures. Over-representation analysis (ORA) was conducted using the gseapy^35^ Python package employing Hypergeometric test and the Hallmark gene sets from MSigDB^36^. For the global analysis, the top 150 proteins ranked by mean SHAP feature importance were used as the input set, with all quantified proteins serving as the background. For risk factor–specific analyses, proteins were ranked by the mean residual SHAP values, and the top 50 proteins for each risk factor were used as input for ORA. P-values were adjusted for multiple testing using the Benjamini-Hochberg false discovery rate procedure.

### Protein expression quantitative trait locus (pQTL) analysis

Protein levels that were used as input for the proteomic aging clock were subjected to pQTL analysis. We used genotypes^37^ imputed with the 1000 genomes reference panel and retained all variants with minor allele frequency greater than 5%. pQTL analysis was performed with tensorQTL. All pQTL with nominal P < 1.0 × 10^−5^ were used as instrumental variables in the Mendelian randomization analysis.

### Mendelian randomization analysis

Mendelian randomization (MR) was performed to assess the potential causal effects of plasma protein levels on aging-related traits.^38–43^ All protein quantitative trait loci (pQTLs; p < 1 × 10^−5^) were selected as instrumental variables (IVs). The weak IVs were firstly filtered by F-statistic > 10. To ensure IV independence, linkage disequilibrium (LD) clumping was performed using PLINK 1.9 (LD *r^2^* < 0.01, window = 1000kb) and the genotypes of the 1000 Genomes reference panel. MR analyses were conducted using the R package, TwoSampleMR. The Wald ratio m ce weighted (IVW) analysis was used as the primary method for proteins with two or more IVs. The significance threshold applied for the primary MR test is nominal P<0.05. When sufficient instruments were available, heterogeneity and directional horizontal pleiotropy were assessed using Cochran’s Q test and the MR-Egger intercept test, respectively, with nominal P < 0.05 indicating significance. For proteins with three or more IVs, if evidence of significant heterogeneity or directional horizontal pleiotropy was detected, three additional sensitivity analyses were performed using the weighted median, weighted mode, and MR-Egger methods, which provide estimates under weaker assumptions.

## Supporting information

All supplementary tables

## Acknowledgements

We thank all participants for their long-term commitment to the KORA study, the staff for data collection and research data management and the members of the KORA Study Group (https://www.helmholtz-munich.de/en/epi/cohort/kora) who are responsible for the design and conduct of the study. We respectfully acknowledge the significant contributions of Dr. Christian Gieger to this project, he passed away before the submission of this manuscript.

## Code availability

All analyses were performed using publicly available python language (version 3.10.16). The code used in this study can be found in the github repository: https://github.com/heiniglab/kora_proteomic_aging_clock

## Data availability

The data are not publicly available because the participants’ consent does not permit it. For research purposes the data can be requested through the KORA Use and Access Hub (https://helmholtz-muenchen.managed-otrs.com/external) subject to the approval by the KORA board.

## Ethics

All study methods were approved by the Ethics Committee of the Bavarian Medical Association (S4 Study: EC No. 99186; F4 study and FF4 study: EC No.06068; Morbidity and Mortality Follow-up in 2021: EC No.20104). All study participants gave written informed consent. The investigations were conducted following the Declaration of Helsinki.

## Supplementary Material / Extended data

**Extended data figure 1:**
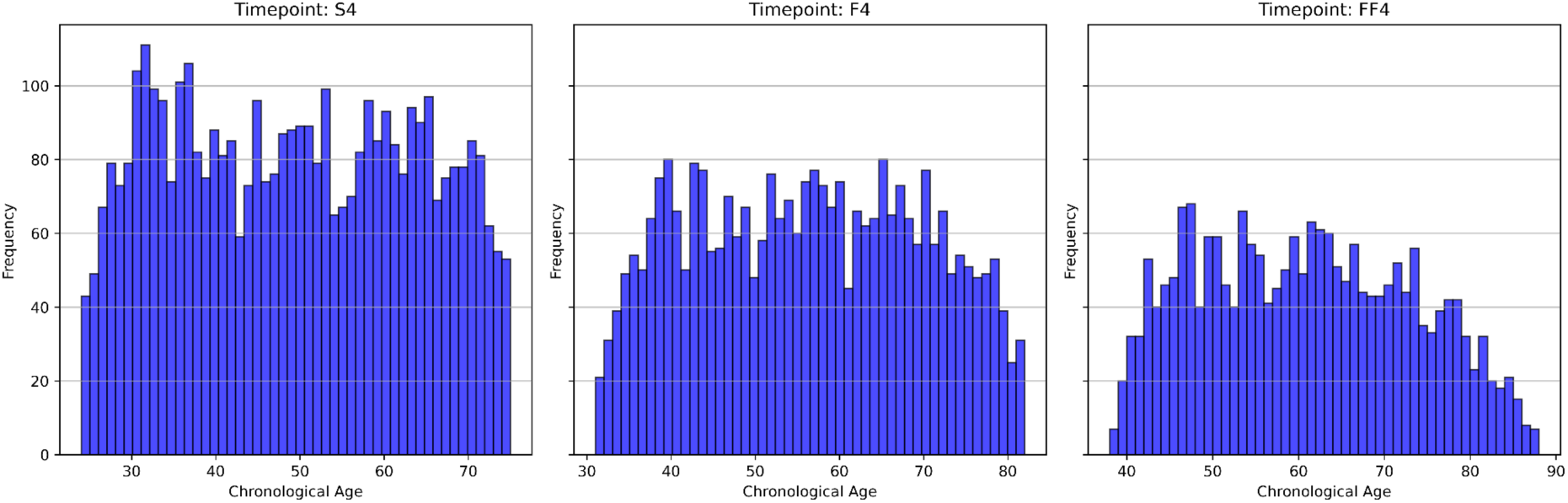
Histograms visualizing age distribution of each timepoint (S4, F4, FF4)

**Extended data figure 2:**
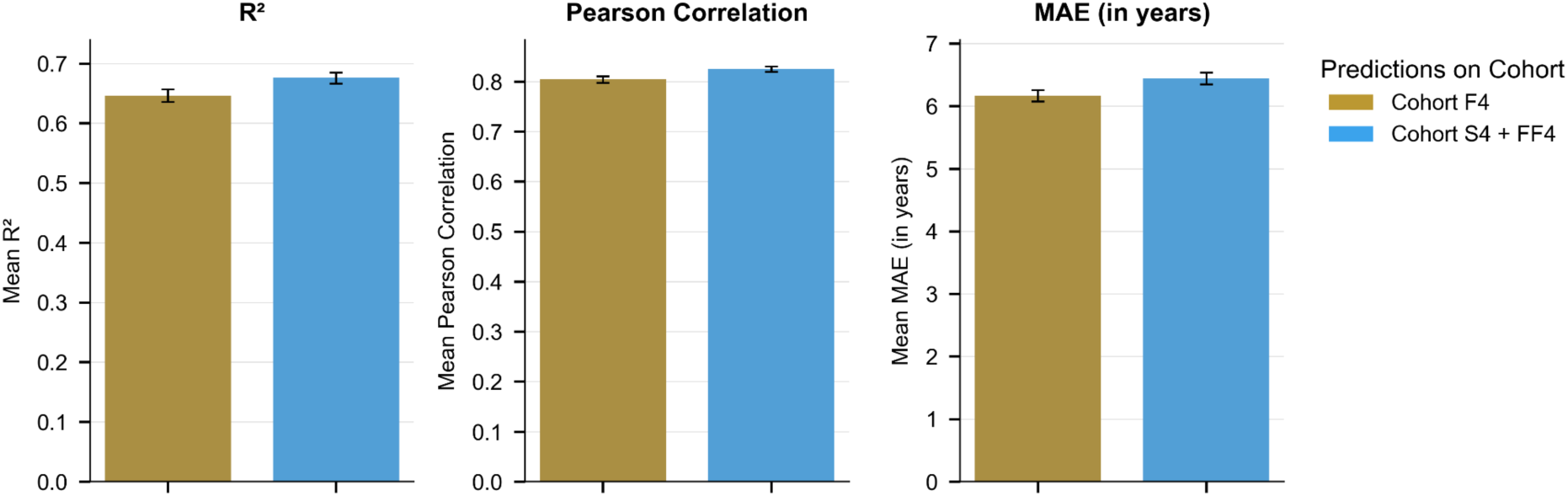
Performance of clocks by mean of R^2^, Pearson correlation and mean absolute error (MAE) fitted on timepoint’s subcohorts S4 and FF4, transferred to F4 subcohort as validation set.

**Extended data figure 3:**
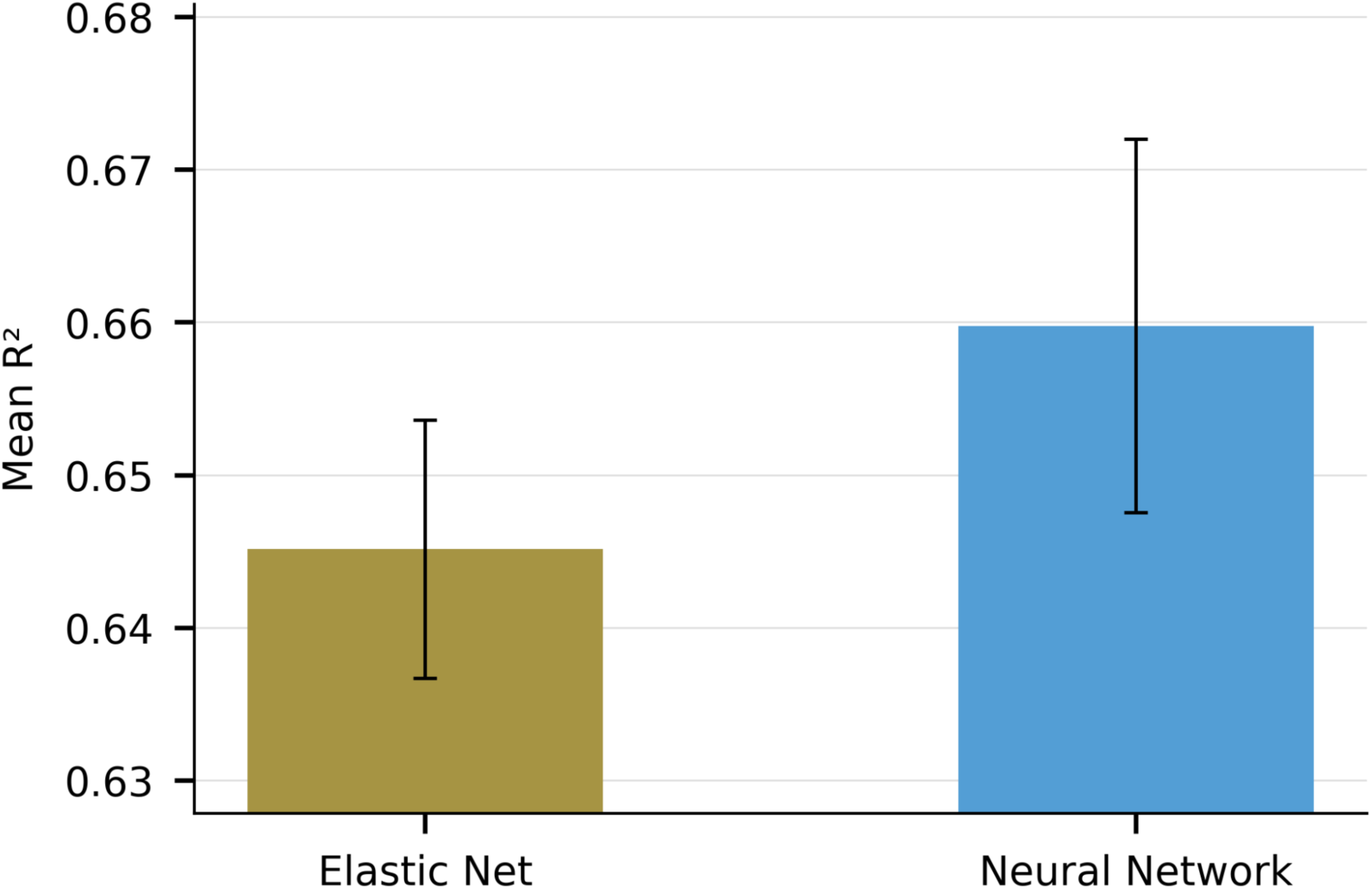
Comparison of the performance of an elastic net model to the neural network model

**Extended data figure 4:**
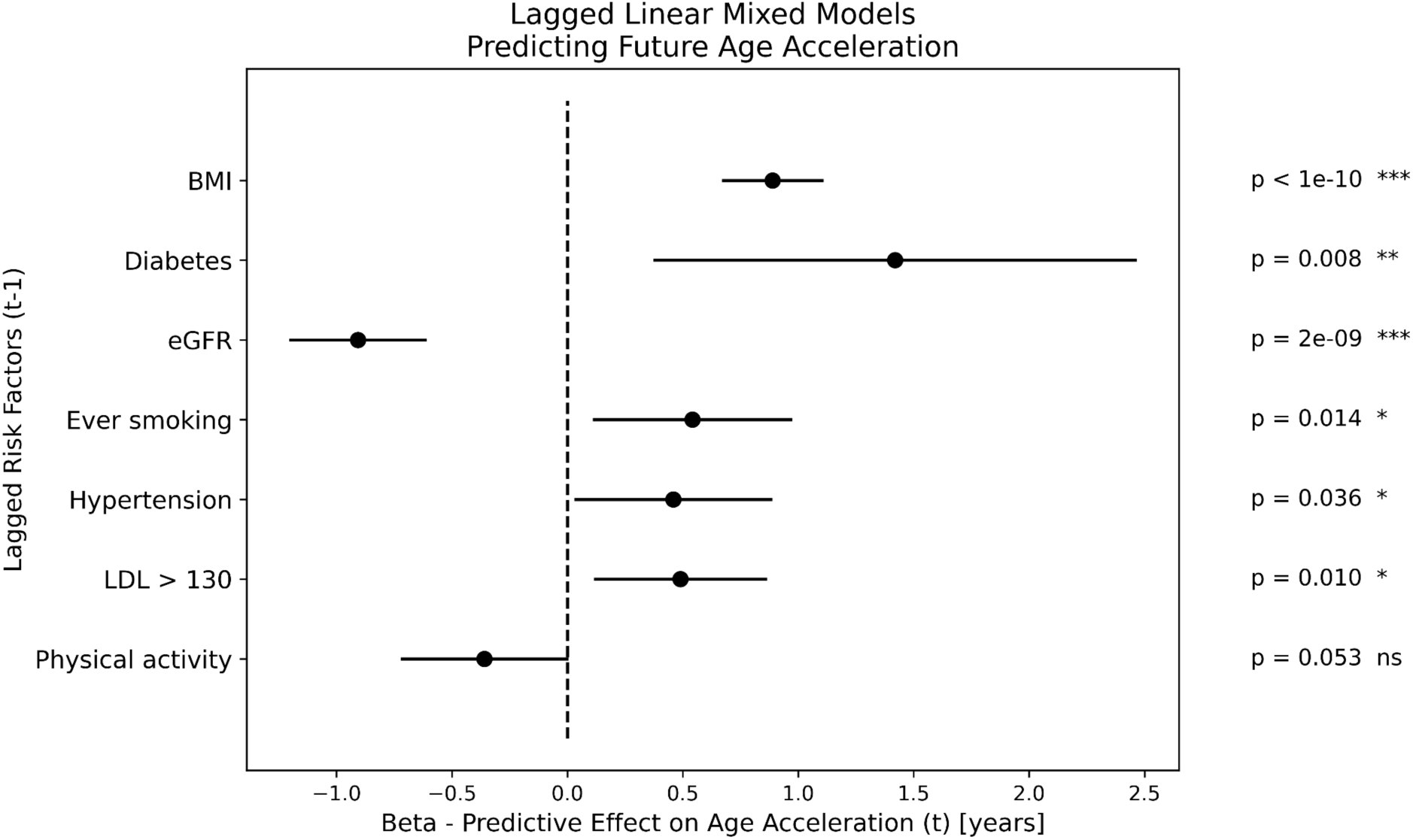
Effect sizes of linear mixed models, based on risk factor exposure from earlier timepoint (t-1, “lagged LMM”). It shows the effect of risk factors from one time point to the age acceleration of later timepoints.

**Extended data figure 5:**
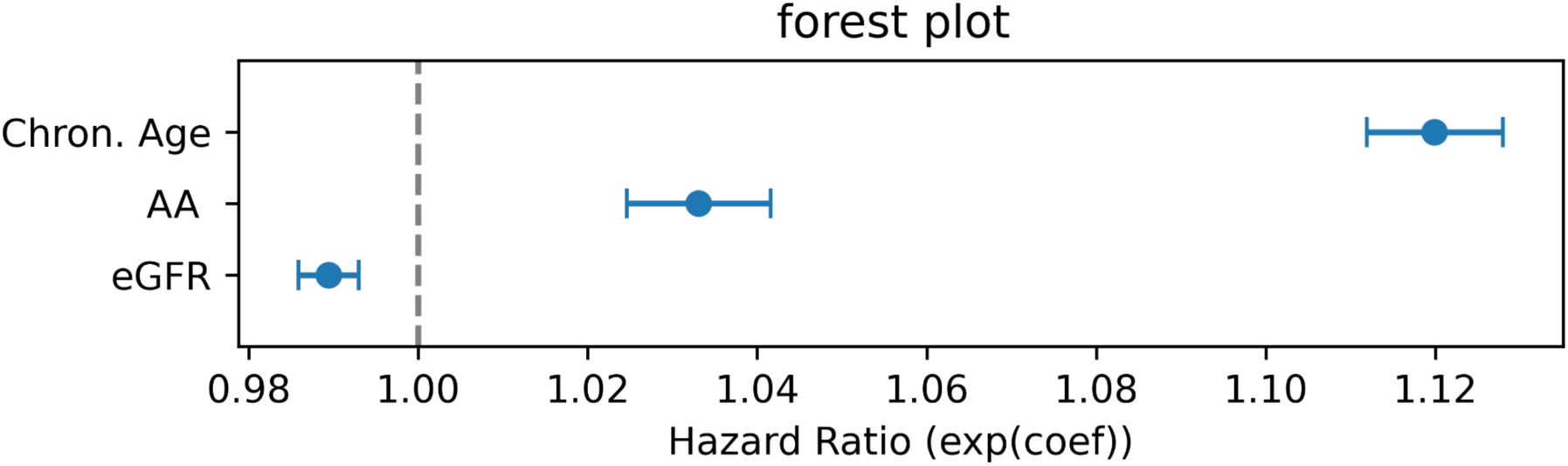
Hazard ratios of the cox model, with chron. age, AA and age acceleration as covariates

